# Flexibility of the neck-linker during docking is pivotal for function of bi-directional kinesin

**DOI:** 10.1101/2020.05.27.118430

**Authors:** Alina Goldstein-Levitin, Kanary Allhuzaeel, Himanshu Pandey, Larisa Gheber

## Abstract

The role of the neck-linker (NL) element in regulating the functions of bi-directional kinesins is unknown. We report that replacing the NL of the bi-directional kinesin-5 Cin8 with sequences from plus-end directed kinesins produces non-functional Cin8 with defective spindle localization and abolished minus-end directionality and microtubule-crosslinking *in vitro*. Mutation of a single glycine in the NL of Cin8 to asparagine (proposed to serve as an N-latch that stabilizes the docked conformation of the NL in the plus-end directed kinesins) causes defects in the functions of Cin8. Strikingly, in a non-functional Cin8 containing the NL of the plus-end directed kinesin-5 Eg5, a single mutation of the N-latch asparagine back to glycine rescues the *in vivo* and *in vitro* defects. Since such replacement eliminates stabilizing interactions between the docked NL and the motor domain, we conclude that flexibility of NL during docking is pivotal for the function of bi-directional kinesin motors.

## Introduction

Kinesin-5 motor proteins perform essential mitotic functions by providing the force that separates the spindle poles apart during spindle assembly, maintenance and elongation [reviewed in ^1–6^]. These motors are homotetramers, with two pairs of catalytic domains located on opposite sides of a central minifilament ^7–10^. This unique structure enables kinesin-5 motors to crosslink the antiparallel microtubules (MTs) of the spindle and to slide them apart by moving in the plus-end direction of the two MTs that they crosslink ^11–13^. Because of the MT architecture within the mitotic spindles, plus-end directed motility of kinesin-5 motors is essential to separate the spindle poles ^1,5,6^. Moreover, since the catalytic domains of kinesin-5 motors are located at the N-terminus, it was previously believed that such N-terminal motors are exclusively plus-end directed. Recently, however, several studies have demonstrated independently that three kinesin-5 motors – the *Saccharomyces cerevisiae* Cin8 and Kip1 and the *Schizosaccharomyces pombe* Cut7 – move processively towards the minus-end of the MTs in single-molecule motility assays and switch directionality under different experimental conditions ^14–21^. The above-described body of work notwithstanding, the molecular mechanism and physiological implications of such bi-directional motility remain elusive.

The majority of kinesin motors share a mechanical structural element of 10-18 amino acids, termed the neck-linker (NL), which connects the two catalytic domains of the kinesin dimers, enabling them to step on the same MT. For N-terminal plus-end directed kinesins, the NL is located at the C-terminal end of the catalytic domain, between the last helix of the catalytic domain (α6) and the neck helix (α7) that is required for dimerization ^22,23^. It has been proposed that during “hand-over-hand” stepping in the plus-end direction, the NL isomerizes between a disordered “undocked” conformation in the presence of ADP and an ordered, motor-domain-bound “docked” conformation in the presence of ATP ^24–28^. It is believed that the docked conformation is stabilized by N-terminal sequences upstream to the motor domain, termed the cover strand (CS), which form a β-sheet with the docked NL ^29–33^. Additional stabilization of the docked NL conformation is provided by an asparagine residue in the middle of the NL that is predicted to serve as a latch (N-latch) holding the docked NL along the core motor domain and is conserved in processive plus-end directed kinesins ^33^. The importance of the isomerization of the NL between the docked and undocked conformations as a force-generating transition in kinesin motors ^24,25,34–36^ is believed to lie in its role in coordinating nucleotide binding and hydrolysis ^36^ and regulating motor velocity, processivity and force production ^37–44^.

During plus-end directed stepping of pairs of motor domains connected via the NL, the transition to, and stabilization of, the docked NL conformation bring the trailing head forward in the plus-end direction and are thus critical for plus-end directed motility. Since bi-directional kinesin motors can move in both plus- and minus-end directions on the MTs, the dynamics of the NL in these kinesins has to allow stepping in both directions. Thus far, however, the role of the NL in regulating the motor functions of bi-directional kinesins has not been reported. To address this issue, we designed a series of NL variants of the bi-directional kinesin-5 Cin8 motor and examined their functions *in vivo* and *in vitro*. We showed that partial replacement of the NL of Cin8 with homologous sequences from the NLs of plus-end directed kinesins resulted in the generation of non-functional Cin8 variants that could neither move in the minus-end direction nor crosslink MTs *in vitro*. In contrast, partial replacement of the NL of Cin8 with NL sequences from the bi-directional kinesin Cut7 produced a partially active Cin8 variant that was minus-end directed *in vitro*, indicating that the NL regulates the directionality and intracellular functions of a bi-directional kinesin.

Strikingly, in the non-functional Cin8 variant containing sequences from the NL of the plus-end directed vertebrate kinesin-5 Eg5, a single replacement of the conserved N-latch asparagine with glycine rescued the majority of the defects of the nonfunctional variant, both *in vivo* and *in vitro*. The replacement of the N-latch asparagine with glycine is predicted to eliminate stabilizing interactions and increase the flexibility of NL during docking ^33^. Moreover, examination of the structure of the bi-directional kinesin-5 Cut7 reveals that in the docked conformation, its homologous N-latch asparagine is not stabilized in the same manner as that in the plus-end directed kinesins, due to differences in residues at a key position within the motor domain. Thus, we propose that flexibility of NL during docking is a common trait of bi-directional kinesin motors and is crucial for their functions.

## Results

To examine the role of the NL in regulating the functionality of Cin8, we generated a series of variants in which some of the amino acids of the Cin8 NL sequence were replaced with homologous sequences from other kinesin motors (Fig. 1A and B). Based on amino acid alignment, we created Cin8 variants containing NL sequences from two exclusively plus-end directed motors, human kinesin 1 KHC (designated here as Cin8_NL_KHC) and *Xenopus laevis* kinesin-5 Eg5 (Cin8_NL_Eg5) or sequences from the bi-directional *S. pombe* kinesin-5 Cut7 (Cin8_NL_Cut7). In addition, we created Cin8 NL variants with single amino acid replacements with amino acids from the above-mentioned kinesin motors (Fig. 1B). These variants were examined in a series of *in vivo* and *in vitro* assays to study the effect of the above-described replacements on the functionality of Cin8.

**Figure 1:**
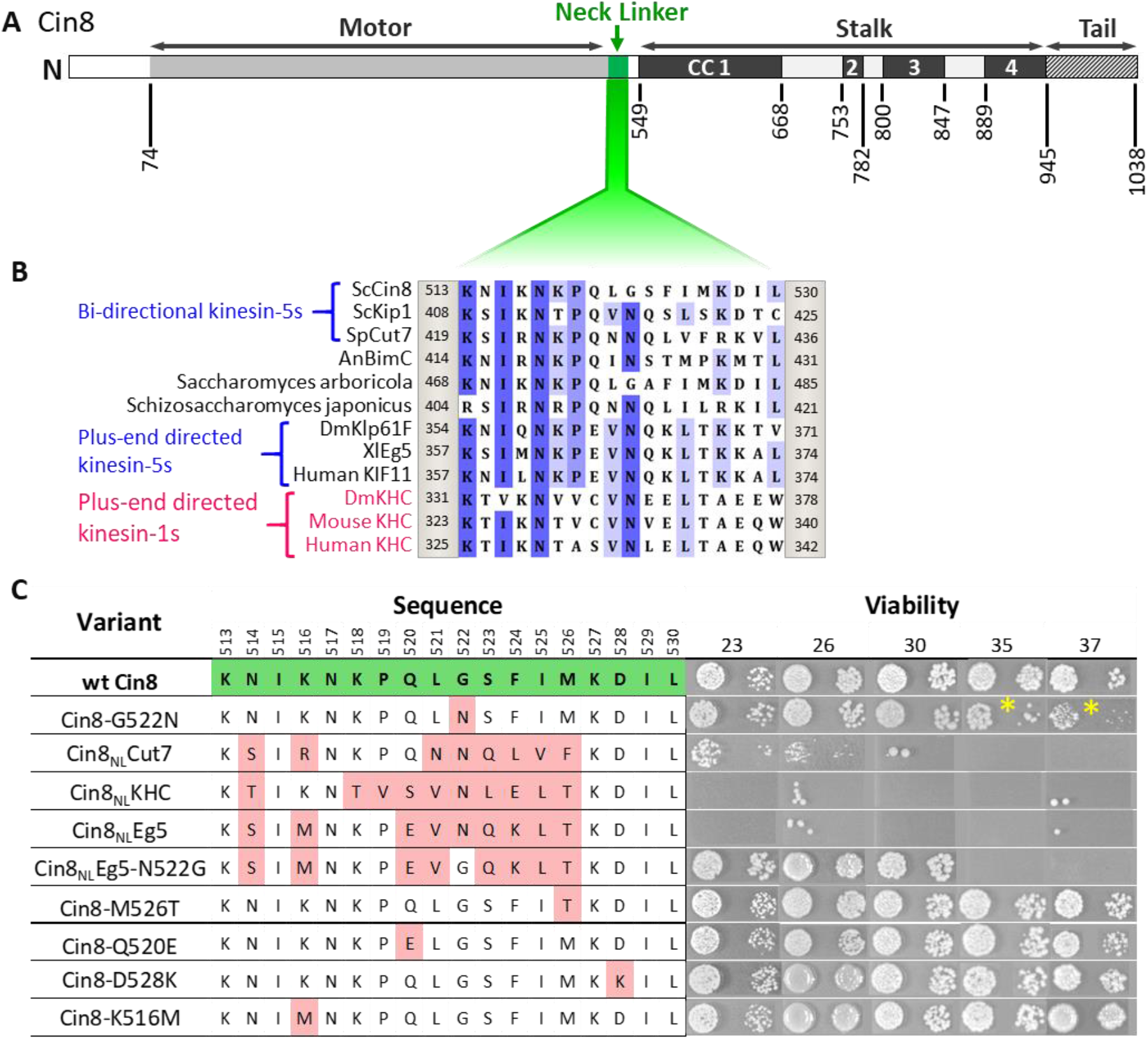
Viability of *Saccharomyces cerevisiae* cells expressing NL variants of Cin8. (**A**) Schematic representation of the Cin8 sequence with amino acid numbers at the bottom flanking the main structural elements of Cin8, indicated on the top; CC: coiled coil. The NL region (green) is expanded below. (**B**) Multiple sequence alignment (MSA) of the NL region of members of the kinesin-5 (black) and kinesin-1 (magenta) families. Known directionality of kinesin motors, i.e., either bi-directional or exclusively plus end directed, is annotated in blue on the left. The positions flanking the presented sequence of each kinesin motor are annotated on the right and on the left of each sequence. The MSA was calculated by the MUSCLE algorithm ^45^ via Unipro UGENE program ^46^. The amino acids are color coded by percentage identity with a 55% threshold. (**C**) Viability of *S. cerevisiae* cells expressing NL variants of Cin8, indicated on the left, as a sole source of kinesin-5. Temperatures (°C) at which cell growth was examined are indicated on the top. Amino acids of the NL are indicated in the middle; positions in the sequence of Cin8 are indicated on the top. The amino acids highlighted in green are those of the wt Cin8 sequence. Amino acids in the NL of Cin8 that were mutated to amino acids from other kinesin motors are highlighted in pink. Asterisks indicate the reduced growth of cells expressing the Cin8-G522N variant at 35°C and 37°C.

### Cin8 containing NL sequences from plus-end directed kinesin motors is not functional in cells

To assess the intracellular functionality of the NL variants, we first examined their ability to support yeast viability as the sole source of kinesin-5 function. Since at least one of the two *S. cerevisiae* kinesin-5 motors, Cin8 and Kip1, is essential for cell viability ^47,48^, the NL variants were examined on a low-copy centromeric plasmid in cells carrying chromosomal deletions of *CIN8* and *KIP1*, following the shuffling-out of the original Cin8 plasmid that covered the double-deletion (Fig. 1C) ^49–51^. We found that NL variants carrying a single amino acid replacement at positions where the consensus of most bi-directional kinesin-5 motors differs from the consensus of the exclusively plus-end directed kinesin-5 motors (variants Cin8-K516M, Cin8-Q520E, Cin8-M526T, and Cin8-D528K) were mostly viable (Fig. 1C), thereby indicating these replacements do not significantly affect the functionality of Cin8 *in vivo*.

In contrast to the single amino acid replacements, Cin8 variants containing partial NL sequences from exclusively plus-end directed kinesin motors (Cin8_NL_KHC and Cin8_NL_Eg5) failed to grow at all the examined temperatures (Fig. 1C), thereby indicating that these variants were not functional in cells. In addition, cells deleted for Cin8 (but with functional Kip1) and expressing the Cin8_NL_Eg5 variant exhibited significantly longer doubling times than cells expressing wild-type (wt) Cin8 (Table 1), as is consistent with the impaired functionality of this variant. Importantly, cells expressing the Cin8 variant with the NL sequence of the bi-directional Cut7 (Cin8_NL_Cut7) were viable at 23⁰C and 26⁰C, but not at temperatures above 30⁰C (Fig. 1C), thereby indicating that at least some of the Cin8 functions are preserved in the Cin8 variant containing the NL sequence from a bi-directional kinesin. Doubling time of cells expressing the Cin8_NL_Cut7 variant was significantly longer than those of cells expressing wt Cin8 in both the presence and the absence of Kip1 (Table 1), which is consistent with the partial functionality of this variant. Moreover, in cells with functional Kip1, variants containing the NL sequence from the plus-end directed Eg5 and bi-directional Cut7 accumulated with monopolar spindles (Fig. 2), indicating that both variants were defective in the spindle assembly function in *S. cerevisiae* cells.

**Figure 2:**
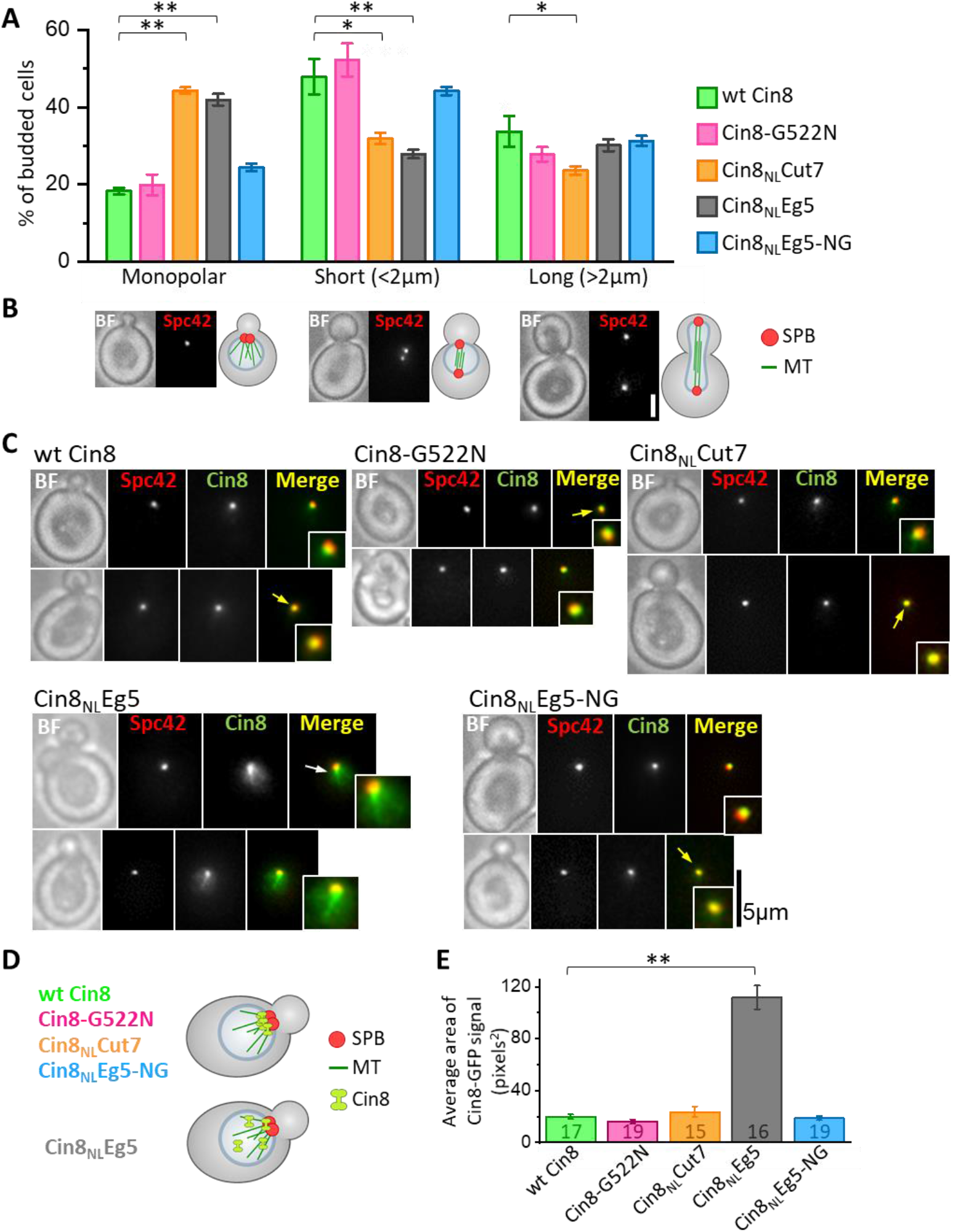
Intracellular phenotypes of NL variants. The examined cells were deleted for the chromosomal copy of *CIN8* (in the presence of *KIP1*) and express tdTomato-tagged SPB component Spc42 and the 3GFP-tagged NL variants. **(A and B)** Spindle length distribution of cells expressing NL variants of Cin8. **(A)** The average percentage of budded cells at the different spindle length categories; (1) Monopolar, (2) Short <2 μm and (3) Long >2 μm (±SEM). NL variants are indicated on the right. In each experiment, at least 200 cells were examined, and spindles were categorized according to their shapes and lengths (see Materials and Methods). For each NL variant, 3-4 experiments were performed. \**p* < 0.05; \*\**p* < 0.01, compared to wt Cin8. **(B)** Live cell images (left) and schematic representation of cells and spindles (right) of each spindle category, as in (A). BF: bright field; Bar: 2 μm. **(C-E)** Localization of NL variants of Cin8 in cells with monopolar spindles. **(C)** Representative images of cells with small buds and monopolar spindles expressing 3GFP-tagged NL variants of Cin8 are indicated on the top of each panel. Cells were imaged in bright field (BF), red (Spc42) and green (Cin8) fluorescence channels. The inserts show a 200% magnification of the localization of Cin8-3GFP. Yellow arrows indicate co-localization of Cin8-3GFP and Spc42-tdTomato, and white arrows indicate localization of Cin8, which is diffusive in the nucleus and is associated with nuclear MTs. Bar: 5 μm. **(D)** Schematic representation of small budded cells with monopolar spindles showing Cin8-3GFP co-localization with the SPBs (top, as in wt Cin8, Cin8-G522N, Cin8_NL_Cut7 and Cin8_NL_Eg5-NG) and diffusive Cin8-3GFP localization in the nucleus and in association with nuclear MTs (bottom, as in Cin8_NL_Eg5). **(E)** Average area of Cin8-3GFP localization in the nucleus (±SEM) of each variant, indicated at the bottom, was calculated by particle analysis function in ImageJ software (see Materials and Methods). Numbers of the examined cells for each variant are indicated in the graph columns. \*\**p* < 0.01, compared to wt Cin8.

**Table 1:** Doubling time of *S. cerevisiae* cells expressing wt and NL variants of Cin8.

|  | *cin8Δkip1Δ*<sup>a,b</sup> | *cin8Δ*<sup>a</sup> |
| --- | --- | --- |
| wt Cin8 | 152 ± 1 (3) | 127 ± 4 (3) |
| Cin8-G522N | 175 ± 2 (3)* | n.d. |
| Cin8<sub>NL</sub>Cut7 | 210 ± 7 (3)** | 143 ± 3 (4)* |
| Cin8<sub>NL</sub>Eg5-NG | 191 ± 6 (4)** | n.d. |
| Cin8<sub>NL</sub>Eg5 | n.d. | 167 ± 4 (4)** |

<sup>a</sup> Values are average doubling times (min) ± SEM. The number of experiments is shown in parentheses. 3HA-tagged Cin8 variants were expressed in the *kip1Δcin8Δ* and *cin8Δ* strains.
<sup>b</sup> In the *cin8Δkip1Δ* strain, NL variants and wt Cin8 were examined following shuffling-out of the parental pMA1208 plasmid (see Materials and Methods).
\* $p < 0.05$ , \*\* $p < 0.01$ , compared to wt Cin8.

To elucidate which function of Cin8 is maintained in the partially viable Cin8_NL_Cut7, but not the non-viable Cin8_NL_Eg5 variant, we visualized the cellular localization of these variants, tagged with 3GFP, in cells containing functional Kip1 and bearing a tdTomato-tagged spindle-pole-body (SPB) protein Spc42 (Spc42-tdTomato) ^16,50^. Prior to bipolar spindle formation, when the two SPBs had not yet separated, wt Cin8 and the Cin8 with the NL sequence from the bi-directional Cut7 concentrated near the SPBs, at the minus-ends of nuclear MTs (Fig. 2C-E). This localization pattern is consistent with the minus-end directed motility of these variants on the nuclear MTs (Fig. 2D) ^52^. In contrast, Cin8 containing the NL sequence from the plus-end directed Eg5 (Cin8_NL_Eg5) exhibited diffusive localization in the nucleus (Fig. 2C-E). Quantitative analysis in cells with pre-assembled spindles does indeed indicate that Cin8_NL_Eg5 occupied a significantly larger area than wt Cin8 and Cin8_NL_Cut7 (see Materials and Methods) (Fig. 2E), suggesting that Cin8_NL_Eg5 exhibits reduced affinity to nuclear MTs. Moreover, in contrast to wt Cin8 and Cin8_NL_Cut7, Cin8_NL_Eg5 appeared to be attached to the nuclear MTs (Fig. 2C, white arrow). Such localization, similar to the previously reported pattern of the tailless Cin8 mutant that had lost its minus-end directionality preference ^20,52^, suggests that the minus-end directed motility is impaired in the Cin8_NL_Eg5 variant, although it is maintained in the partially functional Cin8_NL_Cut7. This is probably the reason for the ability of Cin8_NL_Cut7, but not Cin8_NL_Eg5, to function as a sole source of kinesin-5 function in cells.

In summary, the above experiments indicate that replacement of the Cin8 NL sequence with the sequence from plus-end directed kinesin motors produced variants that are not functional in cells, probably due to decreased affinity to MTs and abolished minus-end directed motility on nuclear MTs prior to spindle assembly. In contrast, replacement of the Cin8 NL sequence with a sequence from the bi-directional Cut7 probably maintained minus-end directed motility, resulting in a partially functional variant of Cin8 that can support cell viability as a sole source of kinesin-5.

### Glycine at N-latch position is crucial for the intracellular functions of Cin8

Cin8 contains a glycine at position 522 in the NL, whereas the majority of plus-end directed kinesin motors contain an asparagine at this position (Fig. 1B), which is predicted to serve as a latch (N-latch) stabilizing the docked NL along the core motor domain ^33^. Importantly, we found that the variant in which this glycine had been replaced with an asparagine, namely, Cin8-G522N, exhibited reduced viability at 35⁰C and 37⁰C, when expressed as the sole source of kinesin-5 (Fig. 1C asterisk). Consistently, *cin8Δ kip1Δ* cells expressing the Cin8-G522N variant exhibited longer doubling times than cells expressing wt Cin8 (Table 1), thereby indicating that the functionality of Cin8 was undermined by the replacement of glycine at position 522 with asparagine. To examine the role of this glycine, we generated a mutant of Cin8 containing the NL sequence of the plus-end directed Eg5, with the asparagine at position 522 mutated back to glycine; this mutant is designated Cin8_NL_Eg5-NG. Strikingly, reinstating the original glycine rescued the non-viable phenotype of the Cin8_NL_Eg5 variant, supporting the viability of *cin8Δ kip1Δ* cells at temperatures up to 30℃ (Fig. 1C). The doubling time of *cin8Δ kip1Δ* cells expressing Cin8_NL_Eg5-NG was longer than that of cells expressing wt Cin8 (Table 1). However, in the presence of Kip1, and in contrast to cells expressing the Cin8_NL_Eg5, no accumulation of monopolar spindles was observed in cells expressing the Cin8_NL_Eg5-NG variant (Fig. 2A and B). Finally, prior to spindle assembly, the Cin8_NL_Eg5-NG variant localized near the SPBs, at the minus-end of the nuclear MTs, similarly to the localization pattern of the wt Cin8 (Fig. 2C-E). This pattern suggests that, in contrast to Cin8_NL_Eg5, minus-end directed motility is restored in the Cin8_NL_Eg5-NG variant, which moves in the minus-end direction on nuclear MTs and concentrates at the SPBs ^13^. Taken together, these results indicate that the presence of glycine at position 522, as in the original sequence of Cin8, rescued the majority of defects of the non-functional Cin8_NL_Eg5 variant. These results further indicate that the glycine at position 522 is critical for the functionality of Cin8 in cells.

### Glycine at the N-latch position regulates the motile properties of Cin8 *in vitro*

To understand the intracellular phenotypes of the NL variants, we examined their motor function *in vitro*. In these assays, we characterized the activity of GFP-tagged full-length NL variants, overexpressed and purified from *S. cerevisiae* cells, on fluorescently labeled GMPCPP-stabilized MTs ^13^. We first compared the MT affinities of the NL variants by determining the average number of MT-attached motors per MT length at equal motor protein concentrations. We found that all the NL variants exhibited significantly lower levels of MT-bound motors than wt Cin8 (Fig. 3A), thereby indicating that NL affects the MT-affinity of Cin8. Moreover, we also found that the Cin8_NL_Eg5 variant, which was unable to support yeast cell viability as a sole source for kinesin-5, exhibited the lowest levels of MT-bound motors, indicating that its affinity to MTs is greatly reduced. The majority of Cin8_NL_Eg5 motors bound to MTs at 140 mM KCl were not motile (not shown). At a lower salt concentration, at which the affinity of the motors to MTs is increased ^19^, Cin8_NL_Eg5 exhibited only bi-directional diffusive motility, with no minus-end directed bias, in contrast to wt Cin8, which exhibited processive minus-end directed motility (Fig. 3B). These results indicate that, consistent with its intracellular localization (Fig. 2C-E), the Cin8_NL_Eg5 variant exhibited very low MT affinity and was unable to move towards the minus-end of the MTs.

**Figure 3:**
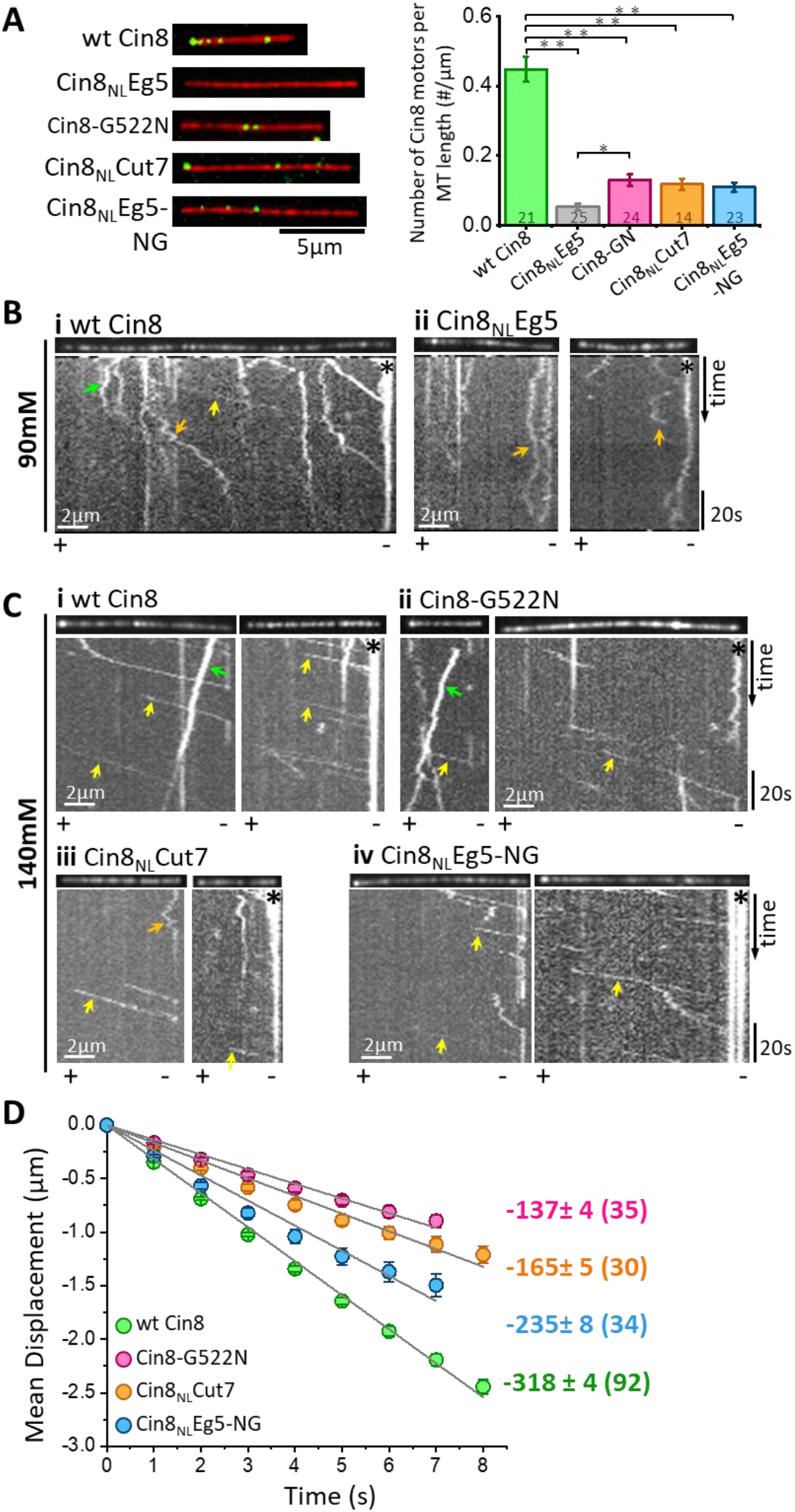
*In vitro* MT binding and single molecule motility assay of NL variants of Cin8. **(A)** MT-binding assay of GFP-tagged NL Cin8 variants in the presence of 1 mM ATP and 140 mM KCl. Left: Representative images of motors (green) bound to fluorescently labeled MTs (red) of the variants; bar: 5 μm. Right: Average number of Cin8 motors per MT length (±SEM). The total number of MT-bound Cin8 motors was divided by the total MT length and averaged over 14-25 observation areas, indicated in the graph columns for each NL variant of 346 μm^2^ (see Materials and Methods); NL variants are indicated at the bottom. \**p* < 0.05; \*\**p* < 0.005. **(B, C)** Representative kymographs of single molecule motility assay of NL variants. **(B)** 90 mM KCl, **(C)** 140 mM KCl. The MTs are shown on the top of the kymographs. The directionality of the MTs, indicated at the bottom of each kymograph, was assigned according to the bright plus-end label and/or by the directionality of fast Cin8 minus-end directed movements ^13,14^. Yellow, orange and green arrows indicate fast minus-end directed, bi-directional, and plus-end directed movements, respectively; asterisks indicate Cin8 clustering at minus-end of MTs. **(D)** Mean displacement (MD) (±SEM) plots of single molecules of Cin8 NL variants as a function of time. The solid lines represent linear fits of the mean displacement (*MD = v·t*, where *v* is the velocity, and *t* is time). Numbers on the right indicate mean velocity (nm/s ± SD), calculated from the linear fits of the MD plots as a function of time. Numbers of analyzed trajectories are indicated in parentheses.

We next examined the motility of the *functional* Cin8 NL variants at a saturating ATP concentration and at a high ionic strength (140 mM KCl). We found that under these conditions, consistent with previous reports ^13,14,52^, all Cin8 NL variants moved processively in the minus-end direction of the MTs (Fig. 3C). We have previously demonstrated that one of the factors that affects the directionality and velocity of Cin8 is its accumulation in clusters on MTs ^13^. Thus, to examine the effects of mutations in the NL sequence on motility but not on motor clustering, we sorted out single molecules of Cin8 from a total population of moving Cin8 particles, based on their intensities (see Materials and Methods). By following the fluorescence intensity of Cin8 particles as a function of time, we observed single events of intensity decrease, of ~45 arbitrary intensity units (a.u.), most probably originating from the photobleaching of single GFP molecules (Fig. S1A of the Expanded Vie section). Since single Cin8 motors are tetramers comprised of four identical subunits ^53^, the maximal fluorescence intensity of a single Cin8 molecule containing four GFP molecules is expected to be ~180 a.u. The intensity distribution of the total Cin8-GFP population was consistent with this notion, in that it exhibited a major intensity peak containing ~65% of Cin8 particles with intensity <180 a.u. (Fig. S1B). The maximal intensity of the peak was ~120 a.u., consistent with an average intensity of single Cin8 molecules containing one, two, three or four fluorescent GFP molecules (Fig. S1B). Based on this analysis (see Materials and Methods), we defined “single Cin8 molecules” as particles of intensity lower than 180 a.u. and analyzed the motile properties only of those Cin8 molecules, thereby ensuring analysis of mainly single molecules of tetrameric Cin8 NL-variants.

We quantified the motility of single molecules of NL variants by tracing their positions on the MTs at each time point, followed by mean displacement (MD) analysis (see Materials and Methods). Consistent with previous reports ^13–15,19^, we found that under high ionic strength conditions, single molecules of wt Cin8 exhibited fast processive movements towards the minus ends of MTs, with a high average velocity of −318±4 nm/s (Fig. 3C and D).

Remarkably, in contrast to the diffusive motility of the Cin8 variant containing the NL sequence of the plus-end directed Eg5 (Cin8_NL_Eg5) (Fig. 3B, orange arrows), mutating the asparagine at position 522 to glycine (Cin8_NL_Eg5-NG) dramatically changed the motile behavior, inducing processive minus-end directed motility (Fig. 3B-D). Although the average velocity of the Cin8_NL_Eg5-NG variant was lower than that of wt Cin8 (Fig. 3D), the processive minus-end directed motility of this variant under the high ionic strength conditions indicates that glycine at position 522 plays a critical role in the interaction with MTs and in stepping in the minus-end direction of the MTs. Consistently, in wt Cin8, mutation of the glycine at this position to asparagine resulted in considerably slower minus-end directed motility, compared with that of the original wt Cin8 (Fig. 3D). Furthermore, Cin8_NL_Cut7, the Cin8 variant with the NL sequence of the bi-directional Cut7 containing asparagine at position 522 (Fig. 1C) also exhibited processive minus-end directed motility, but with reduced velocity compared with that of wt Cin8 (Fig. 3D). These results further support the notion that glycine at position 522 is important for maintaining fast and processive minus-end directed motility of Cin8.

### The NL of Cin8 regulates its MT-crosslinking function

The intracellular functions of kinesin-5 motors are attributed mainly to their ability to crosslink the interpolar MTs of the spindle ^1,5,6^. Thus, to establish additional correlations between intracellular phenotypes and the motor activity of the NL variants, we examined their MT-crosslinking activity *in vitro*. In this assay, purified Cin8 variants were mixed with fluorescently labeled GMPCPP-stabilized MTs in solution, followed by imaging and quantitation of fluorescence intensity of the MTs (see Materials and Methods). Consistent with previous reports ^54^, we found that Cin8 induced the accumulation of MT bundles, thereby increasing the apparent fluorescence intensity of these bundles, compared with that of single MTs (Fig. 4). The MT-bundling activity of wt Cin8 was concentration dependent, inducing MT bundling at a minimal molar ratio of ~1:40 Cin8 motors to tubulin dimer (Fig. 4D).

**Figure 4:**
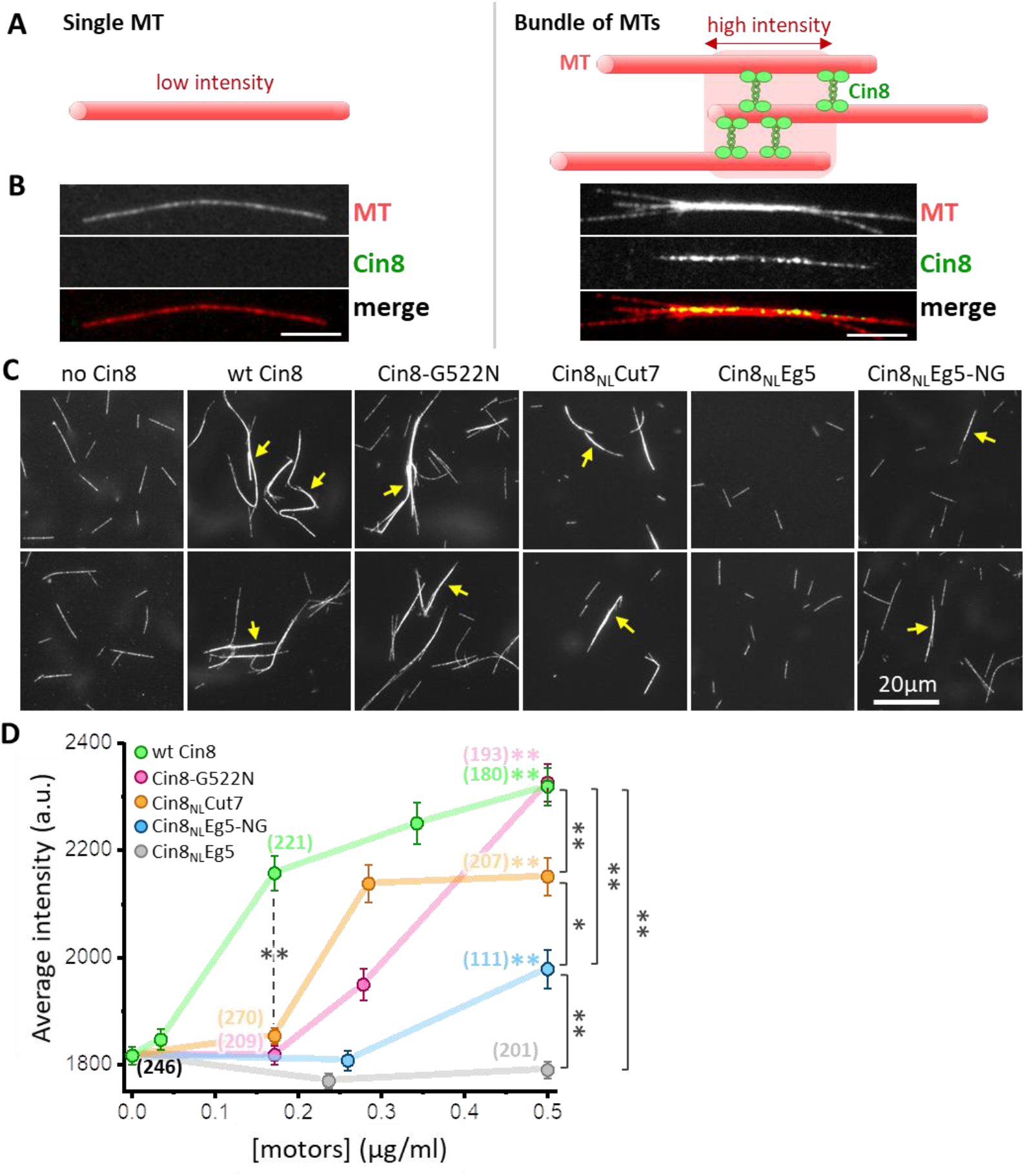
*In vitro* MT bundling by NL variants of Cin8. **(A)** Schematic representation of MTs (red) cross-linked by Cin8 (green). The left panel represents a single MT with a low fluorescence intensity; the right panel represents a high-fluorescence intensity MT-bundle induced by Cin8. **(B)** Representative images of rhodamine-labeled GMPCPP-stabilized MTs (red) in the absence (left) and presence (right) of Cin8-GFP (green). The MT-bundle presented on the right is induced by wt Cin8-GFP, which is co-localized with the bright section of the MT-bundle. **(C)** Representative images of MT-bundles induced by NL variants of Cin8, indicated on the top. Arrows indicate the bright MT bundles. **(D)** Average intensity of MT bundles (±SEM) as a function of the concentration of Cin8 variants, measured by particle analysis using ImageJ (see Materials and Methods). Numbers of particles analyzed for each variant are indicated in parentheses; black: MTs only without motors. Color-coded asterisks for each NL variant indicate comparison to average MT intensity in the absence of motors. \**p* < 0.05; \*\**p* < 0.005.

With the exception of Cin8_NL_Eg5, all NL variants (Fig. 1C) also induced MT bundling in a concentration-dependent manner, with the fluorescence intensity of bundles induced by high concentrations of motors being significantly higher than that of MTs without the addition of motors (Fig. 4D). Moreover, at high motor protein concentrations, the functional NL variants, Cin8-G522N, Cin8_NL_Cut7 and Cin8_NL_Eg5-NG, induced MT bundling to a significantly greater extent, compared with the non-functional Cin8_NL_Eg5. This result indicates that the MT-bundling activity of the NL variants is strongly correlated with the ability of these variants to support cell viability as a sole source for kinesin-5 function (Fig. 1C). However, the *functional* NL variants exhibited reduced ability to bundle MTs, compared with wt Cin8, a finding that is consistent with the reduced affinity to MTs of these variants (Fig. 3A). The Cin8-G255N variant exhibited reduced bundling ability at low and intermediate protein concentrations but reached the MT-bundling levels of wt Cin8 at high protein concentrations. In contrast, both the Cin8_NL_Cut7 and Cin8_NL_Eg5-NG variants exhibited reduced MT bundling compared with wt Cin8 at low and high motor concentrations, with Cin8_NL_Cut7 exhibiting better MT-bundling activity than Cin8_NL_Eg5-NG (Fig. 4D). Finally, in contrast to the Cin8_NL_Eg5 variant which failed to induce MT bundling, a single change of the asparagine at position 522 of Cin8_NL_Eg5 to a glycine, produced a variant that induced MT bundling at high protein concentrations (Fig. 4D), thereby indicating that glycine at position 522 is also important for the MT-crosslinking and bundling activity of Cin8.

In summary, data obtained from the *in vitro* experiments indicates that the NL of Cin8 regulates both its affinity for MTs and the minus-end directed motility of single Cin8 molecules and their ability to crosslink MTs. In addition, the data showed that glycine at position 522, which is equivalent to a conserved asparagine that stabilizes the NL docking in plus-end directed processive kinesins ^33^, is crucial for the fast minus-end directed motility of Cin8 and its ability to crosslink MTs *in vitro*.

## Discussion

The role of the NL dynamics in regulating the motor functions of plus-end directed kinesins has been addressed in a number of studies. However, the current study is the first to demonstrate that mutations in the NL modulate the motile properties and intracellular functions of a bi-directional N-terminal kinesin motor. Data presented here indicates that mutations in the NL regulate MT affinity, minus-end directed motility, and MT crosslinking functions of the bi-directional kinesin-5 Cin8. In turn, these motor functions affect the ability of Cin8 to localize at the spindle poles prior to spindle assembly, to form mitotic spindles, and to support cell viability as a sole source of kinesin-5.

### Minus-end directed motility is important for the intracellular function of a bi-directional kinesin-5

The minus-end directed and switchable directionality of fungal kinesin-5 motors was discovered nearly a decade ago. Although a recent study, based on theoretical simulations, suggests that the minus-end directed motility of the bi-directional *S. pombe* kinesin-5 is necessary for spindle assembly ^55^, experimental support for this notion is still missing. We have recently proposed that in fungi cells, which divide via closed mitosis, minus-end motility of kinesin-5 motors is needed to localize these motors near the spindle poles prior to spindle assembly. At this location, kinesin-5 motors capture and crosslink MTs emanating from the neighboring SPBs and promote SPB separation via antiparallel sliding of the crosslinked MTs ^13^. Analysis of *in vivo* and *in vitro* functions of the NL variants of Cin8 presented here support this model. Our data demonstrates that, compared with wt Cin8, all the examined NL variants exhibited reduced affinity to MTs (Fig. 3A) and an impaired ability to crosslink MTs *in vitro* (Fig. 4C-D). Importantly, our data also shows that all NL variants *that can support the viability of yeast cells* exhibit two common traits: a) they are able to move processively in the minus-end direction *in vitro* (Fig. 3), and b) they localize near the SPBs prior to spindle assembly (Fig. 2C-E), indicating a connection between intracellular functionality, minus-end directed motility, and localization at the SPBs. Differences in minus-end directed velocity per se (Fig. 3C and D) seem to be less important for the intracellular function, as long as the variants can move processively in the minus-end direction. Thus, we conclude that consistent with our model ^5,13^, processive minus-end directed motility that localizes the motors near the SPBs prior to spindle assembly is necessary for mitotic functions of the kinesin-5 Cin8.

### Flexibility of the NL during docking and movement in two directions

The profound difference between the activity of the Cin8_NL_Eg5 and Cin8_NL_Eg5-NG variants indicates that the glycine at position 522 is crucial for the functions of Cin8. The majority of kinesin motors contain asparagine at the position equivalent to glycine 522 of Cin8 (Fig. 1B). It has been suggested that this asparagine plays a central role in the docking of the NL to the catalytic motor domain of the plus-end directed kinesin-1 motors and serves as an N-latch that holds the NL firmly in the motor-domain bound state in the presence of ATP ^33^. Based on the crystal structure of Eg5 in the ATP-like bound state with a docked NL, it appears that this conserved NL asparagine is stabilized by a series of hydrogen bonds with residues within the motor domain (Fig. 5A).

**Figure 5:**
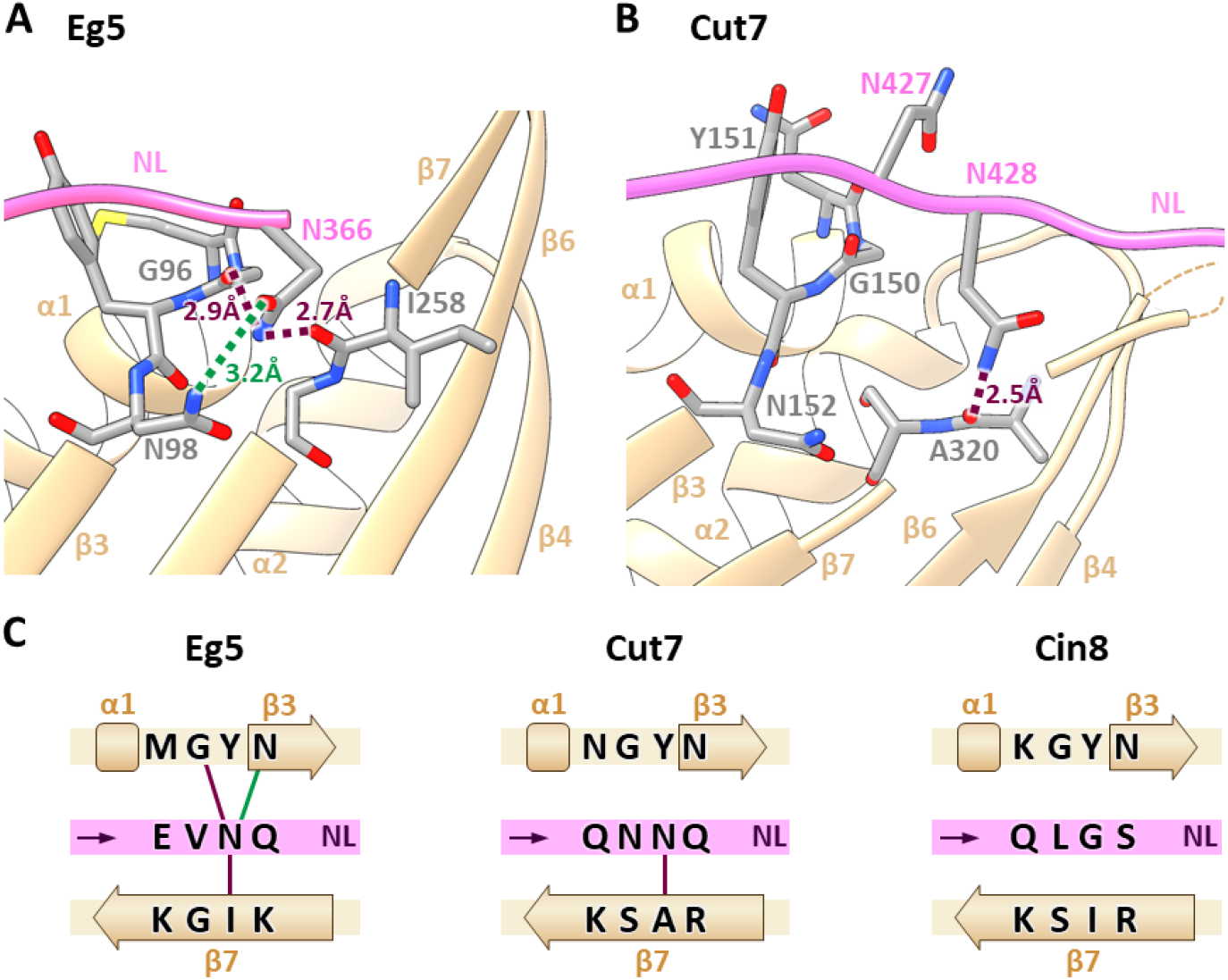
NL docking stabilization of human (Eg5) and *S. pombe* (Cut7) kinesin-5 homologs. **(A)** Crystal structure of the human Eg5 motor domain (light brown) in an ATP-like nucleotide state with NL (pink) docked to the motor domain (PDB:3HQD). Selected amino acids are color coded by element (gray - carbon, red – oxygen, blue – nitrogen, and yellow – sulfur). Nitrogen of asparagine 366 in the NL is in high proximity to the motor domain to enable interaction with the backbone oxygens of isoleucine 258 in β-strand 7 (β7) and glycine 96 in the loop between α-helix 1 (α1) and β-strand 3 (β3) to form hydrogen bonds (dashed purple lines). It is also in suitable proximity to form a hydrogen bond with asparagine 98 in β-strand 3 (β3) (dashed green line). **(B)** Cryo-EM structure of the motor domain of *S. pombe* Cut7 with docked NL in the ATP-like nucleotide state (PDB:5MLV). Color coding of the structural elements as in (A). In the position equivalent to glycine 522 in Cin8, asparagine 428 in the NL of Cut7 can form hydrogen bonds with the backbone oxygen of alanine 320 in β-strand 7 (β7) (dashed purple lines). **(C)** Schematic representation and comparison of the secondary structure and sequence of Eg5, Cut7 and Cin8 (shown in A and B). The secondary structure of the α-helix and β-strands in the motor domain are annotated by squares and arrows, respectively. In each panel, the orientation of the NL, from the N-terminus to the C-terminus, is represented by a purple arrow.

These residues include asparagine 98 in β3 (equivalent to asparagine 159 in Cin8), glycine 96 in the loop between α1 and β3 (equivalent to glycine 157 in Cin8), and isoleucine 258 in β7 (equivalent to isoleucine 414 in Cin8) (Fig. 5A, C and Fig. S2). Plus-end directed kinesin-5 motors are highly homologous at these positions (Fig. S2), indicating that stabilization of NL docking is conserved in these kinesin motors. Although these amino acids are also conserved in the motor domain of Cin8 (Fig. S2), the hydrogen bonds that stabilize NL docking in Eg5 are probably missing (Fig. 5C), due to the presence of glycine rather than asparagine at position 522, which probably results in more flexible NL in the docked conformation in Cin8. In contrast, it is likely that such stabilizing hydrogen bonds are present in the non-functional Cin8_NL_Eg5 variant and in the partially active Cin8_NL_Cut7 and Cin8-G522N variants, which contain asparagine at position 522 (Fig. 1). Replacing this asparagine with the original glycine in the Cin8_NL_Eg5-NG variant probably abolishes these stabilizing hydrogen bonds, thereby partially reinstating the original flexibility of the NL during docking that is required for Cin8 functions. Moreover, based on a recently published CryoEM reconstructed structure, the NL of the bi-directional *S. pombe* kinesin-5 Cut7 is not stabilized by the same interactions as those in Eg5 (Fig. 5B); this difference is probably due to the absence of isoleucine at the position equivalent to I414 in Cin8 (Fig. 5B and C, S2B), indicating that the NL docking of the bi-directional Cut7 is also more flexible than NL docking of the plus-end directed Eg5. Based on these structures, we suggest that flexibility of the NL during docking may be a common trait of bi-directional kinesin-5 motors.

Recent studies indicate that processive stepping in the plus-end direction of kinesin-1 motors is achieved, in part, by a gating mechanism that prevents simultaneous strong binding of the two kinesin-1 motor-domain heads to MTs ^56^. It is believed that this gating is achieved by the coupling between the orientation of the NL, the inter-head tension, and ATP binding and hydrolysis, such that ATP binding in the leading head is prevented until the rear head assumes a weakly bound state or detaches ^27,36,57–61^. However, a different gating mechanism probably takes place during motility of plus-end directed kinesin-5 motors ^44^, which allows for simultaneous strong MT-binding of two motor domains of a dimer in a rigor state with no nucleotides ^62,63^. The two-head strongly bound state has been attributed to the longer and more flexible NL of kinesin-5 motors ^41,59,64^ and may be necessary for force production during antiparallel sliding of MTs in spindle assembly and elongation ^47,65–68^. The bi-directional kinesin-5 motors have the ability to move in both plus-end and minus-end directions, in addition to their ability to crosslink and slide apart antiparallel MTs. Thus, their NL dynamics has to be adapted for bi-directional stepping. Since inducing flexibility by extending the length of the NL was previously shown to enable minus-end directed back-stepping of kinesin-1 dimers ^36^, we propose that increased flexibility during docking of the NL of Cin8, compared to Eg5 (Fig. 5), provides the necessary adaptation to allow bi-directional motility.

### Requirement for flexibility of the NL during docking for the functioning of bi-directional kinesins

In a recent study, the role of NL-docking stabilization in regulating the motility of kinesin-1 was experimentally examined ^42^. That study reported that mutating the N-latch asparagine in the NL of kinesin-1 (equivalent to glycine 522 in Cin8) to alanine, which eliminates stabilizing interactions, increased the plus-end directed motor velocity in a single-molecule motility assay ^42^. Strikingly, we observed the same effect on the velocity of single-molecules of Cin8 during processive movement in the *minus-end direction*. The velocity of Cin8 variants that contain asparagine in position 522 was significantly slower in the minus-end direction than that of wt Cin8 and Cin8_NL_Eg5-NG, which contain glycine at this position (Fig. 3D). These results are consistent with recent CryoEM reports indicating that in the ATP-bound state the NLs of bi-directional kinesin-5 motors assume the docked conformation ^18,29,69^, similarly to kinesin-1 motors ^70,71^. Our results further suggest that stabilization of the NL docking plays similar role in controlling the velocity of plus- and minus-end directed N-terminal kinesin motors.

Although increased flexibility of NL docking similarly affects the velocity of minus-end directed motility of Cin8 and of plus-end directed motility of kinesin-1 [Fig. 3C and D; and ^42^], the intracellular requirements for such flexibility are different for the two types of motor. Whereas for kinesin-1 increased flexibility of the NL reduced force production and impaired the ability of the mutant motors to transport high-load Golgi cargo vesicles ^42^, the intracellular functions of the bi-directional Cin8, reflected in cell viability, localization to the spindle and doubling times, are defective in cells expressing Cin8 variants containing asparagine at position 522 (Fig. 1-3, and Table 1). Furthermore, intracellular functions as well as minus-end directed motility and MT crosslinking were drastically improved by introducing glycine to replace asparagine at position 522 of a non-functional Cin8 variant containing sequences from the NL of Eg5 (Figs. 1-4). Since, as mentioned above, in the presence of glycine, stabilizing interactions of NL docking are considerably reduced (Fig. 5), we conclude that in contrast to plus-end directed kinesin-1, flexibility of NL in the docked conformation is critical for the intracellular functions of bi-directional kinesin-5 motors.

## Materials and methods

### Molecular cloning and strains

NL variants were generated using standard molecular cloning techniques. All mutations were confirmed by sequencing. Plasmids and *S. cerevisiae* strains used in this study are listed in Tables S1 and S2 of the Expanded View section.

### Multiple sequence alignments (MSAs)

MSAs were calculated by the MUSCLE algorithm via Unipro UGENE program ^45^, color coded by percentage identity with a 50-55% threshold. Sequences of the organisms presented in Fig. 1B and Fig. S2 are (from top to bottom): ScCin8 - *Saccharomyces cerevisiae*, Cin8; ScKip1 - *Saccharomyces cerevisiae*, Kip1; SpCu7 - *Schizosaccharomyces pombe*, Cut7; AnBimC - *Aspergillus nidulans*, BimC; kinesin-like protein of *Saccharomyces arboricola*; kinesin-like protein of *Schizosaccharomyces japonicus*; DmKlp61F - *Drosophila melanogaster*, Klp61F; XlEg5 - *Xenopus laevis*, Eg5; human Kif11 – *Homo sapiens* Eg5; DmKHC - *D. melanogaster*, kinesin-1 heavy chain; mouse KHC - *Mus musculus*, kinesin-1 heavy chain; human KHC - *H. sapiens*, kinesin heavy chain isoform 5A.

### Viability assay

The viability of *S. cerevisiae* cells expressing NL variants of Cin8 as the sole source of kinesin-5 was determined as described previously ^13,20,49,51^ (Fig. 1). *S. cerevisiae* strains used for this assay were deleted for their chromosomal copies of *CIN8* and *KIP1* and contained an endogenic recessive cycloheximide resistance gene (*cin8Δkip1Δcyh^r^*), containing a plasmid (pMA1208) encoding for wt Cin8 and a wt dominant cycloheximide sensitivity gene (*CYH*); see Table S2 for the list of *S. cerevisiae* strains. Following transformation with a plasmid encoding for Cin8 NL variants, the pMA1208 plasmid was shuffled out by growth on yeast extract-peptone-dextrose (YPD) medium containing 7.5 μg/mL cycloheximide at 23°C and at elevated temperatures.

### Imaging of *S. cerevisiae* cells

Imaging was performed as previously described ^49,50^ on *S. cerevisiae* cells deleted of the chromosomal copy of *CIN8*, expressing 3GFP-tagged Cin8 NL variants from its native promoter on a centromeric plasmid, and endogenously expressing a SPB component Spc42 with a tdTomato fluorescent protein. Cells were grown overnight and diluted 2 h prior to imaging. Z-stacks of yeast cells were acquired using Zeiss Axiovert 200M microscope controlled by the MicroManager software. Spindle length was measured on 2D projections of the Z-stacks as the distance between the two Spc42-tdTomato SPB components. Cells with monopolar spindles were defined as cells with a small bud and a single Spc42-tdTomato SPB signal. For each variant, three sets of 113-411 cells were categorized according to their SPB morphology and length into three categories: monopolar cells, cells with a short <2 μm spindle, and cells with a long spindle of length > 2 μm. The percentage of budded cells in each category was averaged for three to four experiments for each of the variants, and a statistical analysis was performed by Dunnett’s test ^72^ compared to wt Cin8 at each spindle length category; degrees of freedom (DF) = 10, and F factors 2.89 (α=0.05) and 3.88 (α=0.01). Analysis of Cin8 localization in cells with monopolar spindles (Fig. 2C-E) was performed on 2D projections generated by ImageJ software of monopolar cells. The area of Cin8 localization was measured on images of the fluorescent signal of Cin8-3GFP as follows. First, we generated a mask by thresholding the image using Phansalkar local threshold method of Cin8-3GFP signal in ImageJ. Then, the mask was applied to the original image, resulting an image with zero background. Finally, the area occupied by the Cin8 signal was measured using the particle analysis function in ImageJ. This analysis was performed for 15-19 cells for each Cin8 NL variant and averaged as indicated in the graph columns in Fig. 2E. Statistical analysis was performed by Dunnett’s test ^72^; DF = 92, and F factors 2.55 (α=0.05) and 3.12 (α=0.01).

### Purification of Cin8-GFP

Overexpression and purification of Cin8-GFP from *S. cerevisiae* cells was performed as previously described ^13,19^. Cells expressing Cin8-TEV-GFP-6HIS in a protease-deficient *S. cerevisiae* strain under the GAL1 promoter on a 2μ plasmid were grown in liquid medium supplemented with 2% raffinose. For overexpression, 2% galactose was added for 5 h. Cells were re-suspended in a lysis/binding buffer (50 mM Tris, 30 mM PIPES, 500 mM NaCl, glycerol 10%, 2 mM β-mercaptoethanol, 1 mM phenylmethylsulfonyl fluoride, 1 mM MgCl2, 0.1 mM ATP, 0.2% Triton X-100, Complete Protease Inhibitor (Roche), pH 7.5) and snap-frozen in liquid nitrogen. Cell extracts were prepared by manually grinding in liquid nitrogen in lysis/binding buffer. Ni-NTA beads (Invitrogen) were then incubated with the cell extract for 1.5 h at 4°C, loaded onto a column, and washed with washing buffer (50 mM Tris, 30 mM PIPES, 500 mM NaCl, 30 mM imidazole, 10% glycerol, 1.5 mM β-mercaptoethanol, 0.1 mM Mg-ATP, 0.2% Triton X-100, pH 7.5). Cin8 was eluted with 6 ml of elution buffer (50 mM Tris, 30 mM PIPES, 500 mM NaCl, 250 mM imidazole, 10% glycerol, 1.5 mM β-mercaptoethanol, 0.1 mM Mg-ATP, and 0.2% Triton X-100, pH 7.5). Eluted samples were analyzed by SDS-PAGE and kept frozen in −80°C for further use.

### Single molecule motility assay

The single molecule motility assay was performed on piranha-cleaned salinized coverslips as previously described ^13^: MTs were polymerized by incubating a tubulin mixture containing biotinylated tubulin (T333P), rhodamine-labeled tubulin (TL590M), and unlabeled tubulin (T240) for 1 h at 37⁰C with guanosine-5’-[(α,β)-methyleno]triphosphate (GMPCPP). Then, additional rhodamine-labeled tubulin was added to the polymerizing MTs for 1 h at 37⁰C to form a bright plus-end labeled cap. Flow chambers with immobilized MTs were prepared as previously described ^13^. Cin8, 1.3-0.01 ng/ml, in motility buffer (50 mM Tris, 30 mM PIPES, 110-140 mM KCl, 5% glycerol, 2 mM MgCl_2_, 1 mM EGTA, 30 μg/ml casein, 1 mM DTT, 1 mM ATP, pH 7.2 ATP-regeneration system containing 0.05 mg/mL of phosphocreatine and 0.01 M creatine-kinase) was added to the immobilized MTs and immediately imaged by a Zeiss Axiovert 200M-based microscope, with a 100× objective, equipped with a sCMOS Neo camera. One frame of MTs was captured, followed by time sequence imaging of 90 s with 1-s intervals of Cin8-GFP signal.

### Image and data analysis of the motility assay

Kymographs were generated using ImageJ software for MTs with both ends visible. Directionality of the MTs was assigned on the basis of bright plus-end labeling or by the direction of fast-moving Cin8 particles under high salt conditions ^13,14^. To distinguish between single Cin8-GFP tetramers and clusters, we followed the intensity of stationary Cin8-GFP particles on background-subtracted and corrected for uneven illuminations 90 s with intervals of 1 s by using TrackMate plugin in ImageJ ^73,74^. Since GFP stochastically and irreversibly photobleaches over time, we observed single photobleaching steps, probably representing photobleaching of a single GFP molecule (Fig. S1A). Averaging of the intensity of the photobleaching steps over seven observation fields yielded a value of 45±1 a.u. (SEM) (n = 37), representing the intensity contribution of a single GFP molecule. Therefore, Cin8-GFP particles that have the intensity of four GFPs or less are likely to be single Cin8-GFP tetramers ^74^. MD analysis was performed as previously described ^19,75^. For MD analysis only moving particles with the fluorescence intensity of a single Cin8-GFP molecule (< 180 a.u.) were measured. In addition, measures were taken to minimize the effect of GFP photobleaching on the determination of the Cin8 cluster size. We determined the lifetime of a GFP molecule before photobleaching under our experimental conditions to be 23±3 (SEM) s (n = 40). Consequently, based on this estimation, all the motility measurements were performed only on those Cin8 motors that moved within the first 30 s of each measurement. MD analysis was performed by following the position of such particles, either using TrackMate plugin in ImageJ ^73^ or manually, thereby deducing the displacement at each time interval, followed by averaging the displacement. Average velocity was calculated by fitting the MD analysis to a linear fit corresponding to the equation *M D = ν* · *t*, where ν is the velocity and *t* is time.

### Affinity of Cin8 to MTs

The affinity of Cin8 to MTs was measured on stationary GMPCPP-stabilized fluorescently labeled MTs in the presence of 1 mM ATP and 140 mM KCl, as in the single molecule motility assay, keeping the concentration of Cin8-GFP NL variants constant at ~2.7 ng/ml. One frame of MTs was captured, followed by a one frame of Cin8-GFP signal. Background subtraction was performed by ImageJ software, followed by recognition of Cin8-GFP particles attached to MTs by TrackMate plugin in ImageJ ^73^ in an observation area of 346 μm^2^. For each observation area, the number of MT-bound Cin8-GFP particles was divided by the total length of the MTs and averaged for 14-25 observation areas for each Cin8 NL variant as indicated in the graph columns in Fig. 3A. Statistical analysis was performed by one-way ANOVA all pairwise comparison analysis with Tukey correction; DF = 4, and F factor 61.7.

### MT bundling assay

The MT bundling assay was performed by mixing GMPCPP-stabilized fluorescently labeled MTs, as in the single molecule motility assay, with Cin8-GFP NL variants at concentrations ranging from 0 to 0.5 μg/ml in the presence of 140 mM KCl (and the absence of ATP). The mixture was incubated for 10 min at room temperature and imaged by a Zeiss Axiovert 200M-based microscope, as described for the single molecule motility assay. All images were captured under the same illumination conditions and processed by ImageJ, without any manipulation of images. To calculate the average intensity of MT bundles or single MTs, we first generated a mask by thresholding the image using Phansalkar local threshold method of rhodamine-labeled MTs signal in ImageJ. Then, the mask was applied to the original image, resulting an image with zero background. Finally, average intensity of the MTs or MT-bundles was calculated by particle analysis function in ImageJ, keeping a constant threshold, and averaged for 111-270 MTs/MT bundles for each Cin8-GFP NL variant concentration. Statistical analysis was performed by one-way ANOVA all pairwise analysis with Tukey correction; DF = 3 and F factor 49.1, and DF = 5 and F factor 70.8 for Cin8 concentrations of 0.17 μg/ml and 0.5 μg/ml respectively.

### Doubling time of *S. cerevisiae* cells

The doubling time of *S. cerevisiae* cells was determined as follows: *cin8Δ* or *cin8Δkip1Δcyh^r^* cells were transformed with a centromeric plasmid expressing one of the Cin8 NL variants and grown overnight. In *cin8Δkip1Δcyh^r^* cells, the original plasmid encoding a wt Cin8 and a dominant cycloheximide sensitive gene (CYH) (pMA1208) was shuffled out by addition of 7.5 μg/ml cycloheximide to the growth medium. The cells were diluted to OD_600_ ~0.2 2 h prior to the start of the experiment. The OD_600_ of cells then was measured over time for a culture growing at 26°C, and a plot of 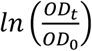 as a function of time was generated, where OD_t_ is the OD_600_ at a given time, and OD_0_ is the OD_600_ at the initial time point. Finally, the doubling time was calculated using the slope of the plot generated according to the equation: 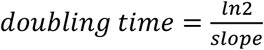. For each variant, the experiment was repeated three to four times. Statistical analysis was performed by Dunnett’s ^72^ test compared to wt Cin8; DF = 10, and F factors 2.67 (α=0.05) and 3.77 (α=0.01), and DF = 9, and F factors 2.76 (α=0.05) and 3.74 (α=0.01) for *cin8Δ* and *cin8Δkip1Δ* respectively.

### Statistical analysis

Data was first examined for normality by the Kolmogorov-Smirnov test ^76^. If normality was rejected, data was subjected to an appropriate Box-Cox transformation ^77^ to yield normal distributions. Next, significant differences were assessed using ANOVA, followed by either all pairwise comparisons with Tukey correction ^78^, or comparisons to wt Cin8 as a control, according to Dunnett’s method ^72^.

### 3D model structural modeling

3D structural models of *H. sapiens* Eg5 (PDB ID: 3HQD) and *S. pombe* Cut7 (PDB: 5MLV) motor domains in an adenylyl imidodiphosphate (AMPPNP)-bound state were depicted by UCSF Chimera software ^79^.

## Acknowledgements

We thank Prof. Levi Gheber, Department of Biotechnology Engineering, BGU, for assistance with image analysis and statistical analysis of the data. We thank Dr. Mary Popov, Ms. Nurit Siegler and Ms. Tatiana Zvagelsky of the L.G. group for critical reading of this manuscript. This research was supported in part by the Israel Science Foundation grant (ISF-386/18) and the United States - Israel Binational Science Foundation grant (BSF-2015851), awarded to L.G.

## Author contributions

A.G.L. Designed and performed experiments, analyzed data and wrote the manuscript; K.A. Performed experiments and analyzed data. H.P. analyzed data and wrote the manuscript; L.G. Conceived the study, designed experiments, analyzed data and wrote the manuscript.

