## Supplemental Information for "Flexibility of the neck-linker during docking is pivotal for function of bi-directional kinesin"

### Figure S1

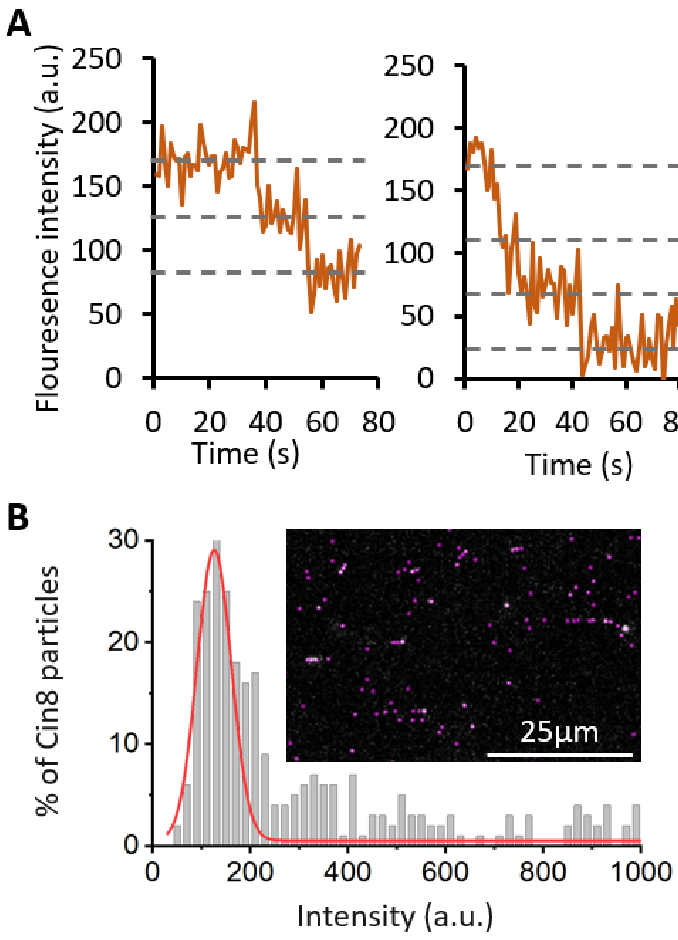


**Figure S1: Fluorescence intensity bleaching and distribution of Cin8 molecules.** (**A**) Fluorescence intensity as a function of time for selected Cin8-GFP particles. Dashed lines represent the fluorescence intensity levels between photobleaching steps. The intensity of the bleaching steps was found to be 45±1 (SEM) a.u. (n = 37). (**B**) Fluorescence intensity distribution histogram of Cin8-GFP particles in the first frame of time-lapse imaging. The red line represents a Gaussian fit of an intensity peak with a center at ~120 a.u. by OriginLab software. This peak contains %64 of the molecules and represents the intensity peak of single Cin8-GFP tetramers. Since the average intensity of a single GFP molecules is ~45 a.u., the average intensity of single Cin8-GFP tetramers is the average fluorescence intensity of one, two, three and four fluorescent GFP molecules, which is ~112 a.u. Detection of particles and their fluorescence intensity determination were performed following background subtraction (see Materials and Methods). The insert is a representative image of field in which the purple circles are particles recognized by the TrackMate plugin in ImageJ (Tinevez et al., 2017).

### Figure S2

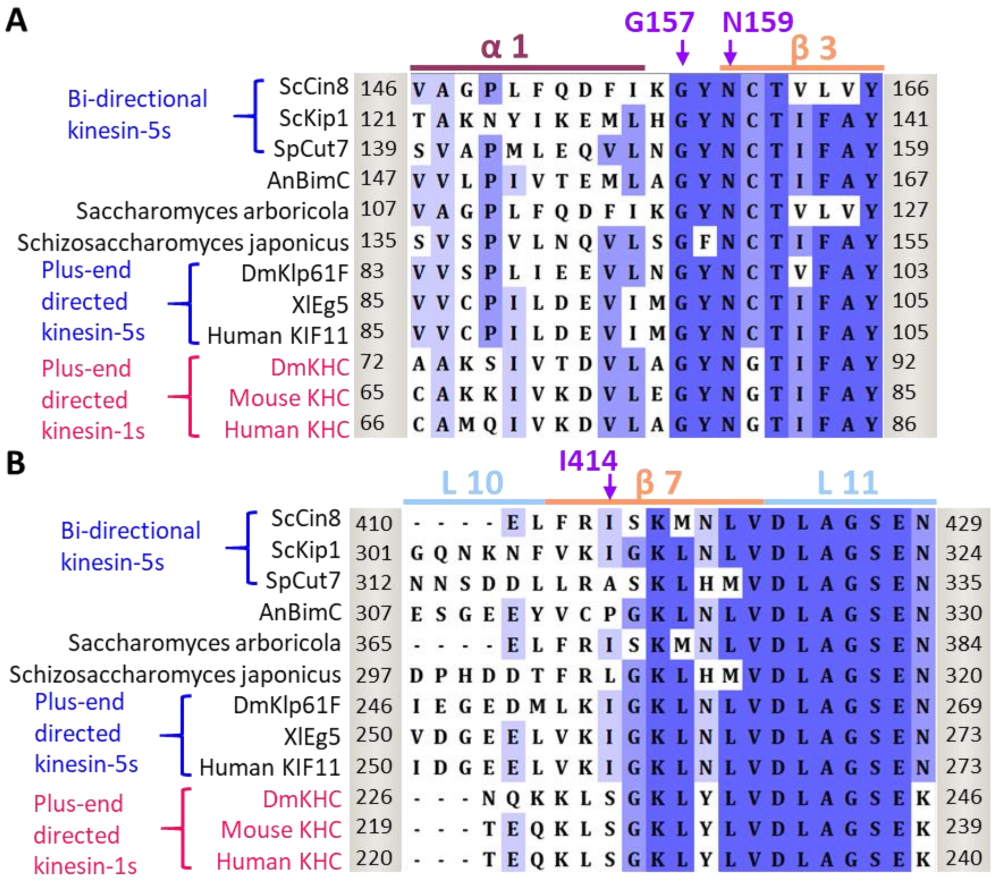


**Figure S2: Multiple sequence alignment (MSA) of regions in the motor domain that probably interact with the NL in docked conformation**. MSA of: **(A)** α-helix 1 (α 1, dark purple), β-strand 3 (β 3, orange), **(B)** loop 10 (L 10, light blue), β-strand 7 (β 7 orange) and loop 11 (L 11, light blue) of members of the kinesin-5 (black) and kinesin-1 (magenta) families. **(A, B)** Positions in the motor domain that are likely to interact with the NL when the NL is docked to the motor domain and to stabilize the docked conformation of the kinesin are shown in purple (glycine 157, asparagine 159 and isoleucine 414 in Cin8). Known directionalities of kinesin motors, i.e., either bi-directional or exclusively plus end directed, are annotated in blue on the left. The positions flanking the presented sequence of each kinesin are annotated on the right and on the left of each sequence. The MSA was calculated by the MUSCLE (Edgar, 2004) algorithm via Unipro UGENE program (Okonechnikov et al., 2012). The amino acids are highlighted by color coding of percentage identity with a 50% threshold. From top to bottom, the presented sequences are: ScCin8 - *Saccharomyces cerevisiae*, Cin8;  ScKip1 -

*Saccharomyces cerevisiae*, Kip1; SpCu7 - *Schizosaccharomyces pombe*, Cut7; AnBimC - *Aspergillus nidulans*, BimC; kinesin-like protein of *Saccharomyces arboricola*; kinesin-like protein of *Schizosaccharomyces japonicus*; DmKlp61F - *Drosophila melanogaster*, Klp61F; XlEg5 – *Xenopus laevis*, Eg5; human Kif11 – *Homo sapiens* Eg5; DmKHC - *D. melanogaster*, kinesin-1 heavy chain; mouse KHC - *Mus musculus*, kinesin-1 heavy chain; human KHC - *H. sapiens*, kinesin heavy chain isoform 5A.

### Table S1: List of plasmids used in this study

| Plasmid | Genotype |
|---|---|
| pMA1208 (Gheber et al., 1999) | *CIN8, CYH2, LEU2, CEN* |
| pVF68 | *CIN8-3EGFP, URA3, CEN* |
| pLG73 | *Cin8$_{NL}$KHC-3EGFP, URA3, CEN* |
| pLG74 | *Cin8$_{NL}$Eg5-3EGFP, URA3, CEN* |
| pLG75 | *Cin8-G522N-3EGFP, URA3, CEN* |
| pLG76 | *Cin8-M526T-3EGFP, URA3, CEN* |
| pLG77 | *Cin8-Q520E-3EGFP, URA3, CEN* |
| pLG80 | *Cin8$_{NL}$Eg5-N522G-3EGFP, URA3, CEN* |
| pLG81 | *Cin8-D528K-3EGFP, URA3, CEN* |
| pLG82 | *Cin8-K516M-3EGFP, URA3, CEN* |
| pLG83 | *Cin8$_{NL}$Cut7-3EGFP, URA3, CEN* |
| pOS7 | *$P_{GAL1}$-CIN8-TEV-EGFP-6His, LEU2, 2µ* |
| pSAG37 | *$P_{GAL1}$Cin8$_{NL}$Eg5-TEV-EGFP-6His, LEU2, 2µ* |
| pSAG38 | *$P_{GAL1}$Cin8-G522N-TEV-EGFP-6His, LEU2, 2µ* |
| pSAG49 | *$P_{GAL1}$Cin8$_{NL}$Eg5-N522G-TEV-EGFP-6His, LEU2, 2µ* |
| pSAG50 | *$P_{GAL1}$Cin8$_{NL}$Cut7-TEV-EGFP-6His, LEU2, 2µ* |
| pRS316 | *URA3, CEN* |
| pKA1 | *CIN8-3HA, URA3, CEN* |
| pKA2 | *Cin8$_{NL}$Eg5-N522G-3HA, URA3, CEN* |
| pKA3 | *Cin8-G522N-3HA, URA3, CEN* |
| pKA4 | *Cin8$_{NL}$Cut7-3HA, URA3, CEN* |
| pKA5 | *Cin8$_{NL}$Eg5-3HA, URA3, CEN* |

#### Table S2: *Saccharomyces cerevisiae* strains used in this study

| Yeast strain | Genotype | Experiment |
|---|---|---|
| LGY 620 | *MATa, ura3-52, leu2-3,112, his3-Δ200, lys2-801, ade2-101, cyh2$^r$, cin8::HIS3, kip1::HIS3, (pMA1208: CIN8, CYH2, LEU2, CEN)* | Cell viability and doubling time |
| LGY 727 | *MATa, ura3-52, leu2-3,112, his3-Δ200, lys2-801, ade2-101. cin8::LEU2* | Doubling time in the presence of Kip1 |
| LGY 1694 | *MATα, ura3-52, leu2-3,112, pep4-3, prb1-1122, reg1-501, gal1* | Overexpression of Cin8 NL variants for motility assays |
| LGY 3989 | *MATa, ura3-52, leu2-3,112, his3-Δ200, lys2-801, ade2-101. cin8::LEU2, SPC42::Spc42-tdTomato, kanMX* | Live cell imaging |
